# Site-1 Protease inhibits mitochondrial metabolism by controlling the TGF-β target gene MSS51

**DOI:** 10.1101/2022.08.24.504591

**Authors:** Muhammad G. Mousa, Lahari Vuppaladhadiam, Meredith O. Kelly, Terri Pietka, Shelby Ek, Karen C. Shen, Gretchen A. Meyer, Brian N. Finck, Rita T. Brookheart

## Abstract

The mitochondrial response to changes in cellular energy demand is necessary for cellular adaptation and organ function. Many genes are essential in orchestrating this response, including the transforming growth factor (TGF)-β1 target gene MSS51, which is an inhibitor of skeletal muscle mitochondrial metabolism. Despite the potential importance of MSS51 in the pathophysiology of obesity and musculoskeletal disease, how MSS51 is regulated is not entirely understood. Site-1 Protease (S1P) is a Golgi-resident protease that is a key activator of several transcription factors required for cellular adaptation. However, the role of S1P in muscle and mitochondrial function are unknown. Here, we identify S1P as a negative regulator of muscle mass and mitochondrial metabolism. Disruption of S1P in mouse skeletal muscle and cultured myofibers leads to a reduction in MSS51 expression, increased muscle mass, and increased mitochondrial oxygen consumption. The effects of S1P deficiency on mitochondrial activity are counteracted by overexpressing MSS51, suggesting that S1P inhibits mitochondrial metabolism by regulating the expression of MSS51. Furthermore, S1P suppression enhances TGF-β signaling via the AKT pathway, potentially explaining muscle hypertrophy in S1P deficient mice. The discovery of S1P as a regulator of mitochondrial metabolism and muscle mass expands our understanding of TGF-β signaling and suggests this protease could be a target for therapeutic intervention in muscle.

## INTRODUCTION

Mitochondria are essential for the cellular response to physiologic and pathologic stimuli. These stimuli elicit dynamic changes in cellular energy demand and substrate availability that require cellular adaptation. Disrupted mitochondrial function can contribute to a failure in adaptation and is associated with several human diseases including muscular dystrophies and sarcopenia – skeletal muscle disorders that are associated with decreased muscle mass and mitochondrial function (1–4). Studies have focused on identifying therapeutic targets to enhance mitochondrial function and thus improve adaptability in human disease states. One key example of this in skeletal muscle is the TGF-β family of proteins that enhance muscle growth and mitochondrial metabolic capacity (5–7). Despite advances in our understanding of the role mitochondria play in adapting to cellular stress elicited by physiologic and pathophysiologic conditions, the molecular mechanisms by which changes in mitochondrial bioenergetics are regulated and, in the case of disease, disrupted, are not yet fully understood.

Site-1 Protease (S1P; also known as subtilisin/kexin-isozyme 1 or PCSK8) coordinates the adaptive response to physiologic or pathologic stimuli through its regulated intramembrane proteolysis (RIP) of key regulators important for maintaining cellular homeostasis (8). S1P-mediated RIP is required for the proteolytic activation of several membrane-bound transcription factors, most notably the sterol regulatory element-binding proteins (SREBP) and ATF6, a key arm of the unfolded protein response (9, 10). Through RIP, S1P coordinates several important signaling pathways associated with human disease and organismal development (e.g., lipid/sterol biosynthesis, lysosomal biogenesis, and the unfolded protein response) (11–20). We previously described a patient with a *de novo*, gain-of-function mutation in S1P who exhibited altered skeletal muscle mitochondrial morphology and myoedema (21). These findings suggested a previously undetected role for S1P in skeletal muscle function and metabolism. Despite the important implications of S1P in human disease and organismal development and its potential influence on skeletal muscle, few studies have directly explored the role of S1P in muscle.

Here, we show that S1P controls mitochondrial metabolism and age-associated muscle mass loss. Although germline deletion of S1P is lethal, in the present study skeletal muscle-specific S1P knockout (S1P^smKO^) was well tolerated and the resulting mice were overtly normal (22). Interestingly, glycolytic muscle fibers from S1P^smKO^ mice show increased maximal mitochondrial respiration and are resistant to age-associated muscle mass loss. Our data suggest that S1P inhibits mitochondrial metabolism by controlling the mitochondrial-resident gene MSS51 and that this regulation partially occurs through the TGF-β1 signaling pathway. These data unveil a previously unknown role for S1P in the regulation of mitochondrial metabolism and muscle mass and identify a potential mechanism by which this occurs.

## RESULTS

### S1P deletion in skeletal muscle increases muscle mass

S1P function has been widely studied in liver and bone, with an emphasis on its role in cellular lipid homeostasis and proteostasis (11, 14–16, 20, 22). We recently described a patient with a gain-of-function mutation in S1P with a pronounced skeletal muscle phenotype; potentially implicating S1P in skeletal muscle function and metabolism (21). To determine the role of S1P function in skeletal muscle, we first examined S1P gene expression levels in various murine muscle groups by quantitative PCR (qPCR). S1P (encoded by the *Mbtps1* gene) is expressed in mouse skeletal muscle (gastrocnemius, tibialis anterior, and soleus), with S1P mRNA levels highest in the gastrocnemius compared to other muscle groups tested (Figure 1A). S1P gastrocnemius mRNA levels were similar to levels in the liver, an organ widely used to study S1P function (20, 22) (Figure 1A).

**Figure 1.**
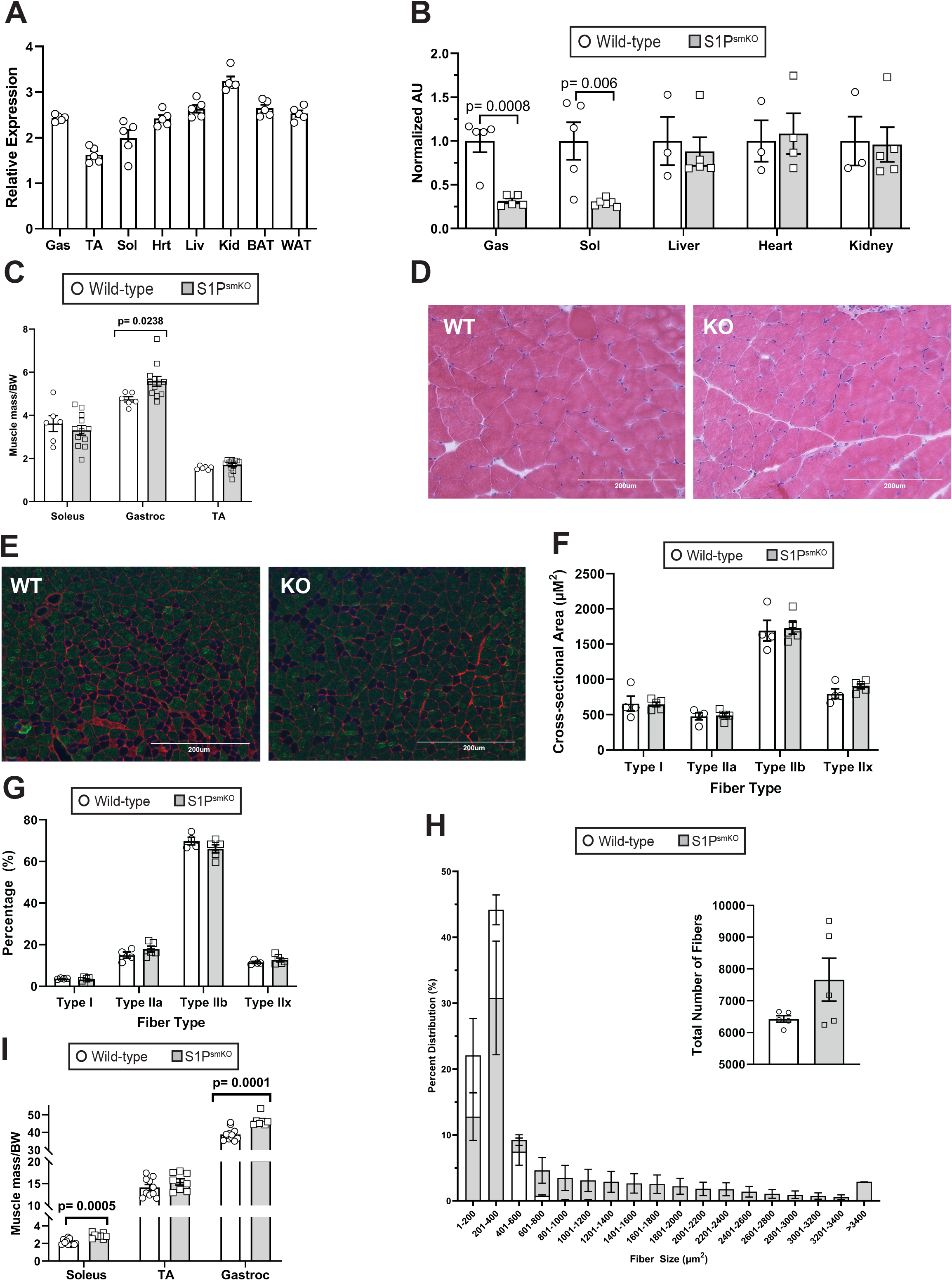
S1P deletion in skeletal muscle increases muscle mass. (A) S1P mRNA expression levels in the indicated mouse organs. n=5 per group. (B) S1P mRNA levels in control (Wild-type) and S1P^smKO^ skeletal muscles and other organs. n=3-6 per group. (C) Normalized muscle mass of soleus, gastrocnemius, and TA of 12-week-old S1P^smKO^ and WT mice. Muscle masses were normalized to body weight (BW). n=6-12 per group. (D-E) H&E and fiber type staining of mid-belly sections of the gastrocnemius of S1P^smKO^ and WT mice. (F-H) Fiber size (cross-sectional area), percent distributions of fiber types, total number of fibers, and fiber size distributions were quantified from mid-belly sections of fiber-type stained images. n=4-5. (I) Normalized muscle mass of soleus, TA, and gastrocnemius of aged S1P^smKO^ and WT mice. Muscle masses were normalized to body weight (BW). n=8-12 per group. Representative images are shown for D and F. Gas, gastrocnemius; TA, tibialis anterior; Sol, soleus; Hrt, heart; Liv, liver; Kid, kidney; BAT, brown adipose tissue; and WAT, white adipose tissue; WT, wild type; KO, knockout.

To investigate the role of S1P in skeletal muscle, we generated skeletal muscle-specific S1P knockout mice (S1P^smKO^) by crossing the established S1P-floxed mouse strain (22) with mice expressing Cre recombinase under the control of the human *ACTA1* promoter (Figure 1B). Although germline deletion of S1P is embryonically lethal, homozygous S1P^smKO^ mice were viable and outwardly normal compared to floxed littermate controls (wild type, WT) (22). Quantification of S1P mRNA in the gastrocnemius and soleus of S1P^smKO^ mice by qPCR showed a robust decrease in S1P mRNA levels compared to WT muscles (Figure 1B).

S1P is a key regulator of SREBPs, which activate a series of target genes required for lipid and sterol biosynthesis (23). Deletion of S1P in mouse liver inhibits SREBP activation, decreasing plasma triglyceride and cholesterol levels, underscoring the need for S1P to maintain lipid/sterol homeostasis (22). Based on these observations, we examined whether S1P^smKO^ mice had altered plasma lipid and cholesterol levels. Compared to WT littermates, S1P^smKO^ mice had normal plasma lipid and cholesterol levels, as well as normal body weight and lean and fat mass (Supplemental Figure 1A-B). We also examined activation of the SREBP pathway in S1P^smKO^ by quantifying expression of SREBP target genes in skeletal muscle by qPCR, and observed no differences in target gene expression between S1P^smKO^ and WT muscles (Supplemental Figure 1C).

To examine the impact of S1P loss on skeletal muscle directly, we performed morphological and histological analyses of S1P^smKO^ and WT muscles. At 12-weeks of age, S1P^smKO^ mice exhibited a 17.6% increase in gastrocnemius mass compared to WT gastrocnemius (Figure 1C). Soleus and tibialis anterior masses were similar between knockout and WT mice. Gross histological examination of gastrocnemius by H&E staining indicated no overt differences in fiber organization in S1P^smKO^ mice relative to WT mice (Figure 1D). We next examined fiber type distribution and size. We observed no differences in fiber size (cross-sectional area) or the overall percentages of type I and type II fibers between knockout and WT muscle (Figure 1E-G). However, fiber size distribution was altered in S1P^smKO^ muscle, where the knockout mice exhibited a greater range of fiber sizes compared to WT mice (Figure 1H). Interestingly, total number of fibers did not vary between knockout and WT muscles (Figure 1H inset). Together these data indicate that S1P^smKO^ mice have increased gastrocnemius mass and increased distribution of fiber sizes.

Skeletal muscle-specific deletion of S1P increased gastrocnemius muscle mass (Figure 1C) and deletion of S1P in bone is associated with increased muscle mass in 40-week-old mice (24). These data suggest S1P may control age-associated muscle mass loss. To investigate this, we examined age-associated muscle mass loss in aged (97-week-old) S1P^smKO^ and WT mice. Aged S1P^smKO^ mice had increased skeletal muscle mass in both soleus and gastrocnemius compared to age-matched WT littermates (Figure 1I). Sirius Red staining of gastrocnemius showed no differences in collagen expansion (i.e., fibrosis) between knockout and control mice (Supplemental Figure 2A). Expression levels of fibrotic markers (*Col1a1, Col3a1*, and fibronectin) were also unchanged between aged S1P^smKO^ and WT controls (Supplemental Figure 2B). These data suggest S1P negatively regulates muscle mass during aging.

### S1P is a negative regulator of mitochondrial metabolism

We previously described a patient with a gain-of-function mutation in S1P that exhibited altered skeletal muscle mitochondrial morphology, implicating a role for S1P in mitochondrial function (21). To examine whether skeletal muscle-specific loss of S1P impacts mitochondrial function, we measured pyruvate-mediated mitochondrial respiration in the gastrocnemius of S1P^smKO^ and WT mice. Because S1P is highly expressed in gastrocnemius and the mass of S1P^smKO^ gastrocnemius is greater than WT mice, we focused on this muscle. The gastrocnemius is composed of glycolytic (fast-twitch) and oxidative (slow-twitch) muscle fibers, which vary in their mitochondrial substrate preferences (25). The ‘white’ gastrocnemius, noted for its opaqueness, is primarily composed of glycolytic fibers, while the ‘red’ gastrocnemius mainly consists of oxidative fibers. Thus, we examined the oxidative capacity of the white and red gastrocnemius separately. Pyruvate-mediated mitochondrial respiration was measured in gastrocnemius fibers permeabilized with saponin. We observed no differences in mitochondrial respiration in the red gastrocnemius between S1P^smKO^ and WT mice (Figure 2A). When we measured mitochondrial respiration in the primarily glycolytic fibers of the white gastrocnemius, Complex I + Complex II respiration and electron transport chain capacity were higher in the white gastrocnemius of S1P^smKO^ mice relative to WT mice (Figure 2B). These findings indicate S1P is a negative regulator of mitochondrial metabolism in glycolytic muscle fibers.

**Figure 2.**
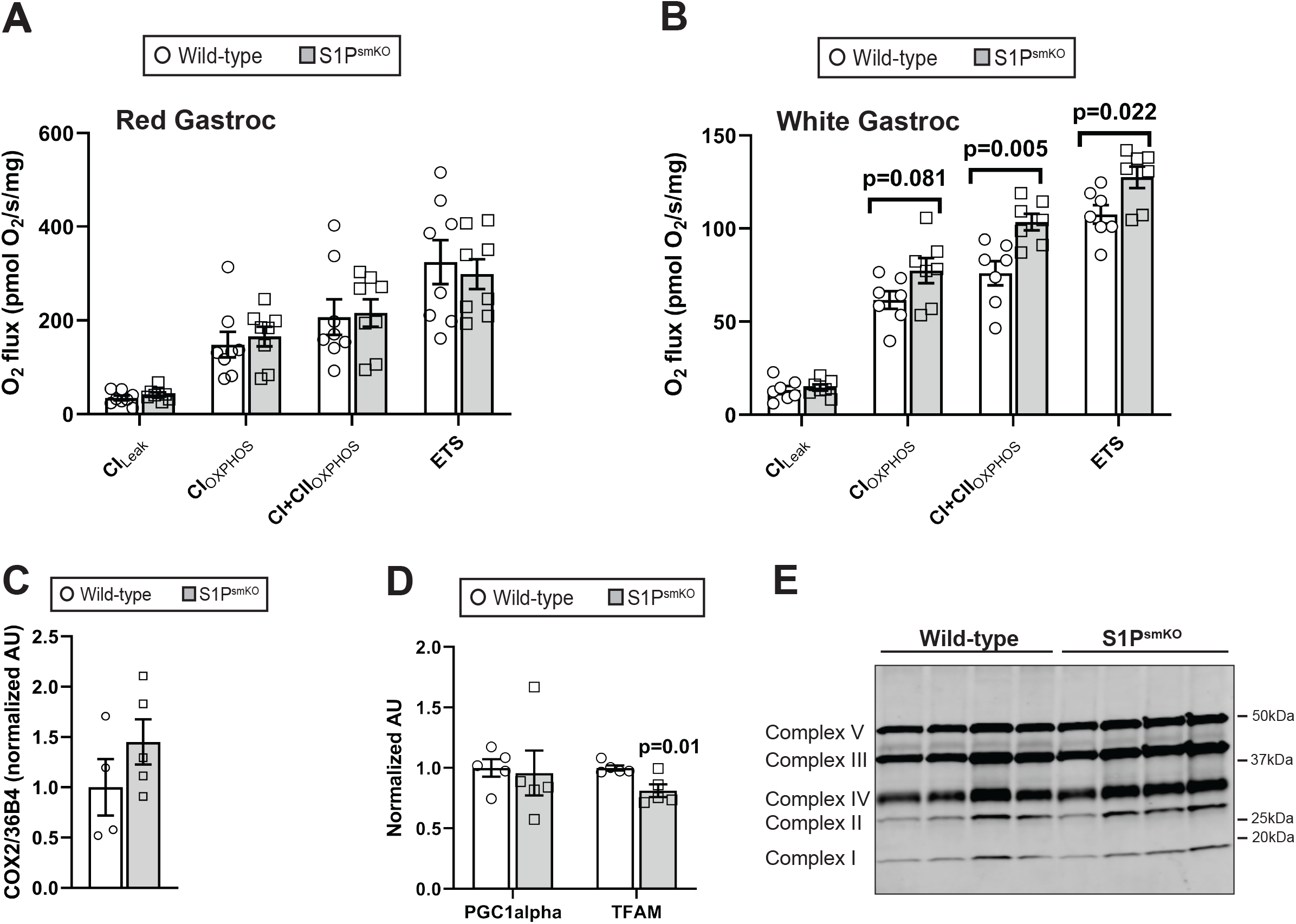
S1P is a negative regulator of mitochondrial metabolism. (A-B) Pyruvate-mediated mitochondrial respiration in red and white gastrocnemius of S1P^smKO^ and WT mice. n=7 per group. (C-D) Mitochondrial DNA content of S1P^smKO^ and WT mice normalized to 36B4. n=4-5 per group. (E) Immunoblotting of oxidative phosphorylation proteins in the gastrocnemius of S1P^smKO^ and WT mice. n=4 per group. Pyr, pyruvate; ETS, electron transport system.

To further characterize the mitochondria of S1P^smKO^ skeletal muscle, we measured mitochondrial DNA content and expression levels of PGC-1α and TFAM in S1P^smKO^ and WT gastrocnemius, markers of mitochondrial number and biogenesis (26). Deletion of S1P from skeletal muscle did not alter mitochondrial DNA content (Figure 2C) and transcript levels of PGC-1α were unchanged between S1P^smKO^ and WT gastrocnemius; however, a small but significant decrease in TFAM transcript levels was observed in S1P^smKO^ muscle compared to WT controls (Figure 2D). To determine whether changes in mitochondrial metabolism were a result of altered expression of mitochondrial electron transport chain (ETC) complexes, we measured ETC protein levels by western blot, and detected no difference in ETC complex protein levels per gram of total protein (Figure 2E). These data suggest that the increase muscle fiber respiration was not due to increased mitochondrial abundance or altered ETC expression levels.

### S1P inhibits mitochondrial metabolism by promoting MSS51 expression

To identify the mechanism by which S1P controls mitochondrial metabolism, we performed RNA Sequencing (RNA-Seq) on RNA isolated from the gastrocnemius of 12-week-old S1P^smKO^ and WT littermates (n=4 per genotype). We identified 75 significantly differentially expressed genes (60 up-regulated and 15 down-regulated) with fold change values greater than 1.5 and p-values greater than 0.05 in the gastrocnemius of S1P^smKO^ mice relative to WT mice (Figure 3A).

**Figure 3.**
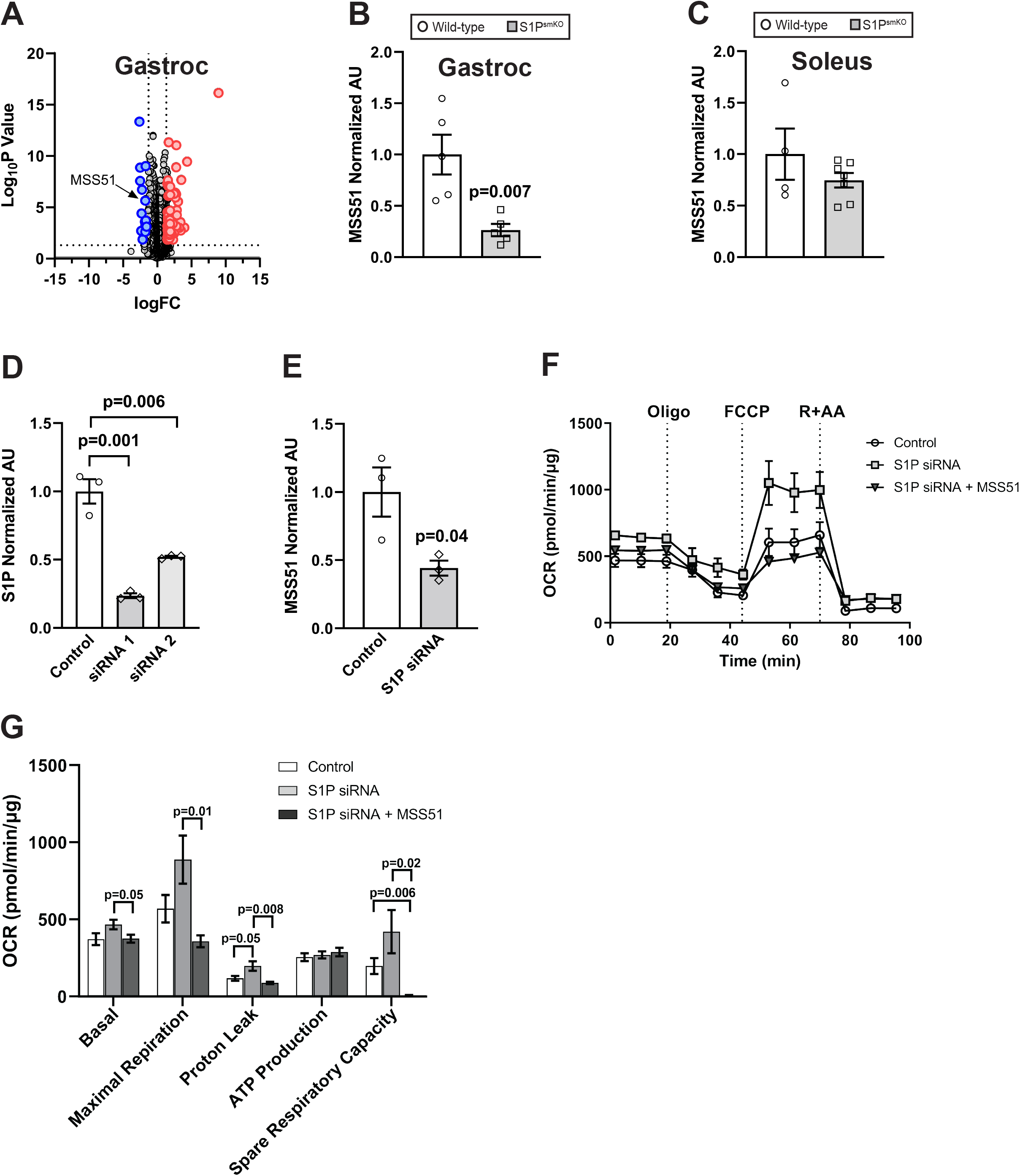
S1P inhibits mitochondrial metabolism by promoting MSS51 expression. (A) Volcano plot of genes identified from RNA-Seq as significantly differentially increased (red dots) and decreased (blue dots) in gastrocnemius of S1P^smKO^ mice relative to WT mice. MSS51 is indicated. n=4 per genotype. (B) qPCR of MSS51 mRNA expression in gastrocnemius of S1P^smKO^ and WT mice. n=5 per group. (C) Knockdown efficiency of custom S1P-targeting siRNAs relative to negative control (control) siRNA in C2C12 cells by qPCR. n=3 per group. (D) qPCR analysis of MSS51 mRNA expression in C2C12 cells transiently transfected with control or S1P-targeting siRNA (S1P siRNA). n=3 per group. (E) Oxygen consumption rate (OCR) of C2C12 cells transiently transfected with control or S1P siRNA plus empty vector or MSS51-Flag tagged plasmid (+MSS51) and quantification of basal, maximal respiration, protein leak, ATP production, and spare respiratory capacity OCR parameters. n=5 per group of a representative experiment of two. Oligo, oligomycin; FCCP, carbonyl cyanide p-trifluoro-methoxyphenyl hydrazone; R+AA, rotenone + antimycin A.

One candidate gene that caught our interest was the protein MSS51, as the expression of MSS51 was dramatically decreased in the gastrocnemius of S1P^smKO^ mice compared to WT mice (Figure 3A). MSS51 is primarily expressed in glycolytic muscle fibers, where it negatively regulates mitochondrial metabolism, but is not involved in mitochondrial biogenesis (6, 7). Indeed, our RNA-Seq analysis and subsequent qPCR analyses of MSS51 expression in the soleus (a primarily oxidative muscle), showed no change in MSS51 transcript levels in S1P ^smKO^ mouse soleus relative to WT mice (Figure 3C). These reported characteristics of MSS51 mirror our observations of muscle group-specific effects of S1P deficiency.

To validate our MSS51 RNA-Seq results, we measured MSS51 mRNA levels by qPCR in the gastrocnemius of S1P^smKO^ and WT mice. Our analysis showed decreased expression of MSS51 transcript levels in the gastrocnemius of S1P^smKO^ mice compared to WT mice, confirming our RNA-Seq results (Figure 3B). To further validate our findings that loss of S1P decreases MSS51 expression, we transiently knocked down S1P in the murine C2C12 cell line using siRNA oligos to target S1P (Figure 3D) and measured MSS51 transcript levels by qPCR. C2C12 cells transfected with scrambled siRNA served as a negative control. Relative to scrambled siRNA cells, depletion of S1P in C2C12 cells decreased MSS51 expression, recapitulating both our RNA-Seq and *in vivo* qPCR results (Figure 3E). These data indicate that S1P is a positive regulator of MSS51 expression.

Because S1P is a positive regulator of MSS51 expression and MSS51 inhibits mitochondrial metabolism (6, 7), we hypothesized that the increase in mitochondrial respiration observed by deleting S1P will be reduced by exogenous MSS51 expression. To test our hypothesis, we used Seahorse respirometry to quantify oxidative respiration in S1P siRNA knockdown cells that over-express either empty vector or FLAG-tagged MSS51. Strikingly, MSS51 expression in S1P knockdown cells led to a decrease in basal oxygen consumption rates, maximal respiration, protein leakage, and spare respiratory capacity compared to S1P knockdown cells expressing empty vector (Figure 3F-G). These data suggest that S1P inhibits mitochondrial respiration by inducing MSS51.

### S1P is involved in the induction of MSS51 expression by TGF-β1

TGF-β1 and its family of ligands induce MSS51 expression through an as yet unknown mechanism (6). To determine if the induction of MSS51 by TGF-β1 requires S1P, we treated S1P knockdown and scrambled (control) C2C12 cells with either vehicle or recombinant TGF-β1 and measured MSS51 expression by qPCR. In the presence of TGF-β1, MSS51 expression was increased in both scrambled and S1P-depleted cells; however, the TGF-β1-mediated induction of MSS51 expression was significantly attenuated in S1P-depleted cells compared to scrambled control treated cells (Figure 4A). These data demonstrate that S1P positively regulates MSS51 expression by TGF-β1.

**Figure 4.**
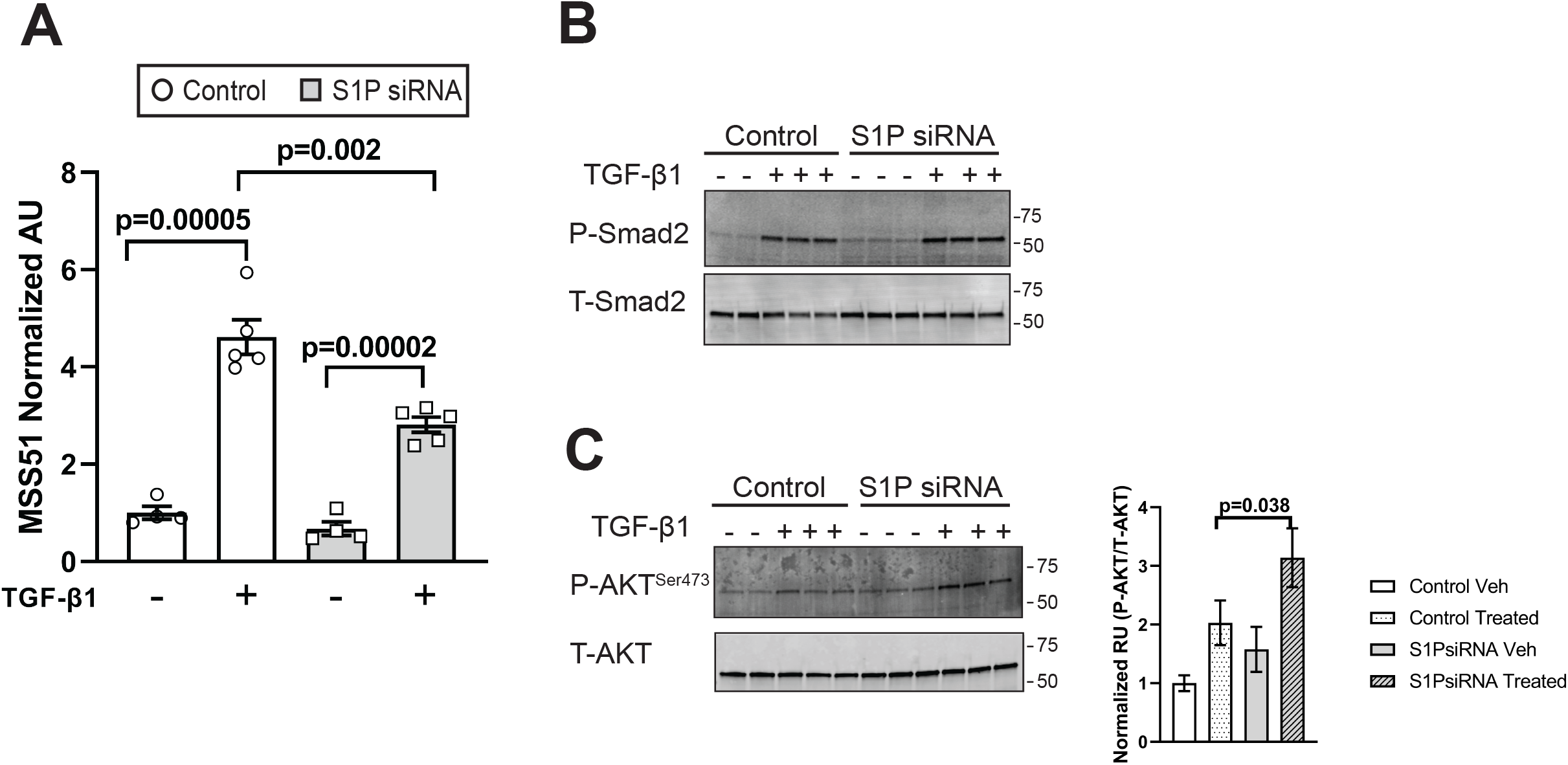
S1P controls MSS51 expression through TGF-β1. (A) MSS51 mRNA expression of scrambled (control) and S1P siRNA transfected C2C12 cells treated with vehicle (-) or TGF-β1 (+) for 5 h after 3 days in differentiation media. n=4-5 per group. (B-C) Western blot of phosphorylated Smad2 (P-Smad 2) and total Smad 2 (T-Smad 2); and phosphorylated AKT (P-AKT^Ser473^) and total AKT (T-AKT) in scrambled (control) and S1P siRNA transfected C2C12 cells treated as in panel (A). n=2-3 per group.

To date, S1P has not been shown to modulate TGF-β1 signaling. TGF-β1 can regulate cellular function through both Smad-dependent and Smad-independent signaling pathways (27). To examine the function of S1P on TGF-β1 signaling and gain insight into the mechanism of S1P-driven MSS51 expression, we first investigated TGF-β1-induced Smad activation. Binding of TGF-β1 to its receptors triggers the phosphorylation and activation of the transcription factor Smad 2 (28), thus Smad 2 phosphorylation is a positive marker of TGF-β1 Smad-dependent signaling. We assessed the phosphorylation status of Smad 2 in whole-cell lysates from vehicle (untreated) and TGF-β1-treated scrambled and S1P-depleted cells by western blotting. Smad 2 phosphorylation was not detectable in untreated cells but Smad 2 phosphorylation was equally induced in scrambled and S1P-depleted cells treated with TGF-β1 (Figure 4B). These data suggest S1P controls TGF-β1-induced MSS51 expression independently of Smad 2 phosphorylation/activation.

We next examined TGF-β1 Smad-independent signaling, which includes TGF-β1-driven AKT activation through phosphorylation of AKT on serine 473 (27). Interestingly, TGF-β1-driven AKT activation is known to antagonize Smad transcriptional activity (27). We assessed levels of phosphorylated AKT^Ser473^ and total AKT in untreated and TGF-β1-treated scrambled and S1P-depleted cells. S1P-depleted cells treated with TGF-β1 had increased phosphorylated AKT compared to treated control cells (Figure 4C). These data suggest S1P depletion leads to increased TGF-β1-dependent activation of AKT, which could also explain observed effects of S1P deletion on muscle hypertrophy.

## DISCUSSION

S1P is a key coordinator of the adaptive response to physiologic and pathologic stimuli. S1P initiates the cleavage and subsequent activation of several regulators required to maintain and restore cellular homeostasis (9–11). Much work has focused on the role of S1P in liver and bone, and its involvement in lipid/sterol homeostasis and proteostasis (9, 10, 13–16, 20, 22). Our previous work suggested a role for S1P in skeletal muscle function and metabolism (21). To date, very little is known about the impact of S1P function on skeletal muscle and whether non-canonical functions for S1P exist in this tissue.

In the present study, we investigated the biological role of S1P in skeletal muscle using skeletal muscle-specific S1P knockout mouse line and identified S1P as a regulator of muscle mass and mitochondrial metabolism. Specifically, S1P^smKO^ mice have increased gastrocnemius muscle mass and, as mice age, this increase in mass is present in both gastrocnemius and soleus muscles relative to age-matched control littermates, implicating a role for S1P in age-associated muscle mass loss. S1P^smKO^ mice also have increased Complex I + Complex II respiration and elevated maximal (+FCCP) respiration. Increased maximal respiration was also recapitulated in S1P siRNA knockdown C2C12 cells. Levels of the mitochondrial-resident gene MSS51 were decreased in both S1P^smKO^ gastrocnemius and S1P-depleted C2C12 cells. Exogenous expression of MSS51 in S1P knockdown cells obliterated the increases in maximal respiration observed, indicating S1P inhibits mitochondrial metabolism by driving MSS51 expression.

Our S1P^smKO^ studies show increased mitochondrial respiration in predominately glycolytic muscle fibers, but not in oxidative fibers of the gastrocnemius. This may be due to increased abundance or activity of S1P in glycolytic fibers relative to oxidative fibers, as has been reported for the S1P substrate SREBP-1c (29). Moreover, MSS51 abundance is concentrated in glycolytic muscle relative to oxidative muscle (6). Indeed, our RNA-Seq and qPCR analysis of mouse soleus, which is primarily composed of oxidative fibers, showed no significant change in MSS51 transcript levels between S1P^smKO^ and control solei. Thus, suggesting S1P-dependent control of mitochondrial metabolism is focused on glycolytic fibers and that S1P may have an as yet unknown function in predominantly oxidative muscle types.

We show that MSS51 expression is decreased in the absence of S1P. In mammals, little is known about the factors that control MSS51 expression, and even less is understood about how MSS51 modulates mitochondrial respiration. Members of the TGF-β1 family of ligands (e.g., TGF-β1 and myostatin) induce MSS51 expression via an as yet unknown mechanism (6). Here, we begin to explore how S1P controls MSS51 expression and show that siRNA depletion of S1P in culture partially inhibits TGF-β1-driven MSS51 expression. One drawback to our siRNA system is that S1P was depleted, not completely deleted and thus it is possible the remaining amounts of S1P enzyme were sufficient to drive blunted, yet detectable levels of MSS51 expression.

Because depletion of S1P impacted TGF-β1-dependent MSS51 expression, this suggested a role for S1P in controlling TGF-β1 signaling pathways. To begin to explore this possibility, we examined whether S1P modulated TGF-β1 signaling via Smad-dependent or Smad-independent signaling pathways and demonstrated that depletion of S1P did not impact Smad 2 phosphorylation, but did increase TGF-β1-induced AKT phosphorylation. Whether S1P controls Smad activity downstream of Smad 2 phosphorylation (i.e, localization and/or activation of Smads) is not known. AKT antagonizes Smad-mediated transcription, thus the increased AKT activation in our S1P siRNA studies suggests AKT may negatively control S1P-driven MSS51 expression by inhibiting Smad activity (27).

Disrupted mitochondrial function and metabolism are associated with Duchenne muscular dystrophy (DMD) (1, 2, 30). In mouse models of DMD (mdx mice), disrupted mitochondrial metabolism was observed early on in disease progression, suggesting disrupted mitochondrial metabolism may contribute to DMD pathophysiology (2). Deletion of MSS51 in mdx mice improves basal and maximal respiration relative to control mdx mice (30). Given our evidence that S1P promotes MSS51 expression, a role for S1P in DMD disease progression is possible.

In addition to controlling mitochondrial metabolism, TGF-β1 family ligands also control muscle mass (5, 6, 31). We observed increased muscle mass in the gastrocnemius of 12-week-old S1P^smKO^ mice, as well as in other muscle types of aged S1P^smKO^ mice. Moreover, S1P^smKO^ mice exhibit increased fiber sizes compared to WT mice, suggestive of muscle hypertrophy. Because AKT controls cell size and AKT activation is increased in S1P-depleted cells, it is possible that the increased skeletal muscle size of our S1P^smKO^ mice may be a result of enhanced AKT activity. These data combined with our observations that S1P regulates MSS51, suggest a possible role for S1P in bridging TGF-β1-dependent control of muscle mass and mitochondrial metabolism. Since S1P inhibitors are in clinical development, inhibition of S1P as a therapeutic target to increase muscle mass in aging or other conditions associated with sarcopenia could be feasible.

In conclusion, these studies identify S1P as a regulator of mitochondrial metabolism and age-associated muscle mass loss. Our data also shed light on the regulation of MSS51 by linking S1P to TGF-β1 signaling. Together, our findings uncover a previously unknown function for S1P in mitochondrial biology and implicate S1P in the adaptation to disruptions in skeletal muscle mass and metabolism. Current work is focused on examining the mechanism(s) by which S1P modulates mitochondrial function and the implications this may have on human disease states and aging.

## ACKNOWLEDGEMENTS

We thank Drs. Jay Horton at UT Southwestern and Linda Sandell at Washington University School of Medicine for the S1P floxed mouse strain. We also thank the Genome Technology Access Center in the Department of Genetics at Washington University School of Medicine for help with genomic analysis. The Center is partially supported by NCI Cancer Center Support Grant P30 CA91842 to the Siteman Cancer Center and by ICTS/CTSA Grant UL1TR002345 from the National Center for Research Resources (NCRR), a component of the National Institutes of Health (NIH), and NIH Roadmap for Medical Research. This publication is solely the responsibility of the authors and does not necessarily represent the official view of NCRR or NIH.

Research reported in this publication was supported by DRC grant P30 DK020579, Washington University Institute of Clinical and Translational Sciences grant UL1TR002345 from the NIH National Center for Advancing Translational Sciences, Nutrition Obesity Research Center NIH grant P30 DK056341, Washington University Musculoskeletal Research Center NIH grant P30 AR057235, and NIH grant K01 HL145326. The content is solely the responsibility of the authors and does not necessarily represent the official views of the National Institutes of Health.

## AUTHOR CONTRIBUTIONS

Conceptualization, R.T.B.; Methodology, R.T.B., B.N.F., G.M.; Formal analysis, R.T.B., T.P., K.C.S., M.O.K.; Investigation, L.V., M.G.M, R.T.B, T.P., S.E., K.C.S., M.O.K.; Resources, G.M., B.N.F.; Writing – Original: R.T.B.; Writing – Review & Editing: R.T.B., B.N.F., G.M., T.P.; Funding Acquisition: R.T.B.; Supervision: R.T.B., G.M.

## DECLARATION OF INTERESTS

Rita Brookheart and Brian Finck are inventors on a pending U.S. non-provisional patent application titled, Methods and Compositions for Improving Exercise Endurance, Performance, or Tolerance (Application Number: 16/732,740). Rita Brookheart holds a provisional patent related to this work (provisional patent application no. 63/370,712).

## METHODS

### Animal Studies

All mouse studies were approved by the Institutional Animal Care and Use Committee of Washington University. Mice were maintained on a standard laboratory chow diet and group housed on a 12 h light/dark cycle. For experiments, 10–97-week-old mice were used. For blood chemistry analyses, chow-fed mice were fasted for 4 h (09:00-13:00h) followed by tail-vein blood withdrawal. Blood was collected by venipuncture of the inferior vena cava and processed for plasma collection via centrifugation in EDTA-coated tubes, and frozen in liquid nitrogen. Skeletal muscle and other organs were harvested and either immediately snap frozen in liquid nitrogen for downstream gene expression analyses or processed for mitochondrial respiration studies or histology.

### Generation of S1P^smKO^ mice (C57BL/6J background)

S1P floxed mice in the C57BL/6J background were previously described (22) and obtained from Linda Sandell at Washington University, with generous permission from Jay Horton of University of Texas Southwestern. HSA-Cre79 mice were obtained from Jackson Laboratory (B6.Cg-Tg(ACTA1-cre)79Jme/J; Stock No. 006149) in the C57BL/6J background. S1P floxed mice were crossed with HSA-Cre79 mice to generate skeletal muscle-specific S1P knockout mice. Littermates not expressing Cre recombinase were used as controls for all experiments. Mice were genotyped for the presence of Cre recombinase and floxed S1P allele (22) using gene-specific primers. Primer sequences are listed in Supplemental Table 1.

### Body composition

ECHO MRI was used to measure body composition in unanesthetized mice using an ECHOMRI 3-1 (ECHO Medical Systems).

### Serum metabolites

Plasma triglyceride and cholesterol levels were measured enzymatically via the Infinity triglyceride (TR22421) and cholesterol (TR13421) assay kits (Thermo Fisher) as per manufacturer’s instructions.

### Histological analyses

Tragacanth gum was placed on top of corks and fresh muscles were vertically placed in the gum so that 1/4 of the muscle was embedded. Samples were submerged in cold (-150 °C) isopentane as described (32) for 20 s, and immediately stored at -80 °C until sectioning. Frozen muscles were transversely cryosectioned into 10 µM thick sections at the mid-belly on a cryostat (Leica Biosystems). Sections were stained with haematoxylin (H&E), Sirius Red or immunostained against myosin heavy chain isoforms (type I (BA-F8), type IIa (SC-71), type IIx, and type IIb (BF-F3); Developmental Studies Hybridoma Bank) and laminin (ab11575, Abcam). Cross sectional area, fiber size, and fiber type distribution of each fiber type was quantified from immunostained fiber type images as reported previously (33).

### Gene expression analysis

Total RNA was isolated from C2C12 cells, skeletal muscle, heart, adipose, kidney, and liver with RNA STAT-60 (Tel-Test Inc) as per manufacturer’s instructions. For tissues, RNA was isolated by disrupting tissue in RNA STAT-60 using 5mM steel beads (Qiagen) and a TissueLyser II (Qiagen). RNA was reverse transcribed into cDNA using the High-Capacity cDNA Reverse Transcription Kit (Applied Biosystems). Quantitative real-time PCR was performed using Power SYBR green (Applied Biosystems) and transcripts quantified on an ABI QuantiStudio 3 sequence detection system (Applied Biosystems). Data was normalized to 36B4 expression, unless otherwise noted, and results analyzed using the 2^-ΔΔCt^ method and reported as relative units to controls. Primer sequences are listed in Supplemental Table 1.

### RNA-Seq Analysis

Gastrocnemius and soleus were harvested from 12-week-old male floxed (wild type) and S1P skeletal muscle-specific knockout mice and immediately snap frozen in liquid nitrogen for a total of n=4 mice per genotype examined. RNA was isolated from tissue as described above. RNA was DNase I treated as per manufacturer’s instructions (RNase-Free DNase Set, Qiagen) then cleaned up and eluted with RNAse and DNAse free molecular grade water (RNeasy MinElute Cleanup Kit, Qiagen), followed by quantification (NanoDrop, ThermoFischer Scientific). RNA with RIN values greater than 8 were accepted for RNASeq. Samples were prepared according to library kit manufacturer’s protocol, indexed, pooled, and sequenced on an Illumina HiSeq. Basecalls and demultiplexing were performed with Illumina’s bcl2fastq software and a custom python demultiplexing program with a maximum of one mismatch in the indexing read. RNA-seq reads were aligned to the Ensembl release 76 primary assembly with STAR version 2.5.1a (34). Gene counts were derived from the number of uniquely aligned unambiguous reads by Subread:featureCount version 1.4.6-p5 (35). Isoform expression of known Ensembl transcripts were estimated with Salmon version 0.8.2 (36). Sequencing performance was assessed for the total number of aligned reads, total number of uniquely aligned reads, and features detected. The ribosomal fraction, known junction saturation, and read distribution over known gene models were quantified with RSeQC version 2.6.2 (37).

All gene counts were then imported into the R/Bioconductor package EdgeR (38) and TMM normalization size factors were calculated to adjust samples for differences in library size. Ribosomal genes and genes not expressed in the smallest group size minus one sample greater than one count-per-million were excluded from further analysis. The TMM size factors and the matrix of counts were then imported into the R/Bioconductor package Limma (39). Weighted likelihood based on the observed mean-variance relationship of every gene and sample were then calculated for all samples with the voomWithQualityWeights (40). The performance of all genes was assessed with plots of the residual standard deviation of every gene to their average log-count with a robustly fitted trend line of the residuals. Differential expression analysis was then performed to analyze differences between conditions and the results were filtered for only those genes with Benjamini-Hochberg false-discovery rate adjusted p-values less than or equal to 0.05.

### Cell culture

All cells were grown at 37 °C with 5 % CO^2^. C2C12 cells (ATCC) were grown in DMEM supplemented with 10 % fetal bovine serum and 1 % pen-strep. To differentiate C2C12 myoblasts into myotubes, cells were grown to 80% confluency, washed with 1x PBS and grown in DMEM supplemented with 2 % horse serum for 2-3 days as indicated in the methods.

### siRNA studies

C2C12 cells were plated onto 6-well plates at a 2 x10^5^ density and 24 h later transfected with either negative control siRNA (Negative Control No.1 siRNA, Life Technologies) or custom siRNAs targeting S1P (Silencer Select siRNAs, Life Technologies) using Lipofectamine RNAiMAX as per manufacturer’s instructions. After 48 h, cells were either harvested for gene expression analysis or differentiated with DMEM supplemented with 2% horse serum. Two days after differentiation, cells were harvested for gene expression analysis. For TGF-β1 treatment studies, three days post-differentiation, cells were treated with 50ng/ml TGF-β1 (R&D) for 5 h then harvested for downstream endpoints.

### Seahorse OCR analysis

Cellular respiration was measured on a Seahorse XFe24 Analyzer (Agilent). C2C12 cells were plated onto 24-well Seahorse XF24 cell culture microplates at a 8,000 cell density and 24 h later co-transfected with either negative control siRNA (Negative Control No.1 siRNA, Life Technologies) with empty vector (pCMV6-Entry; Origene PS100001) or a custom siRNA targeting S1P (Silencer Select siRNAs, Life Technologies) with empty vector (pCMV6-Entry; Origene PS100001) or mouse MSS51-Myc-FLAG (mouse cDNA clone; Origene MR217897) using Lipofectamine 2000 as per manufacturer’s instructions. After 48 h, cells were washed with 1x PBS and switched to differentiation media (DMEM and 2% horse serum) for 3 days. Cells were fed Seahorse XF-DMEM with 1mM pyruvate, 10mM glucose, and 2mM glutamine and incubated in a CO^2^-free incubator at 37 °C for 1 h. Basal oxygen consumption rates were measured first followed by OCR measurements upon sequential addition of oligomycin (1 µM), carbonyl cyanide p-trifluoro-methoxyphenyl hydrazone (FCCP; 1 µM), and rotenone and actinomycin A (0.5 µM each) as per the Seahorse Mitochondrial Stress Test protocol. After completion of the assay, whole protein lysates for each well were quantified by BCA assay and total protein amounts were used for normalization of Seahorse data using Wave Software (Agilent). Basal OCR, maximal respiration, protein leakage, and spare respirometry capacity were calculated using the Seahorse XF Cell Mito Stress Test Report Generator via Wave Software (Agilent) normalized to total protein levels.

### Preparation of permeabilized muscle fibers and high-resolution respirometry

Freshly isolated red and white gastrocnemius sections were immersed in cold BIOPS (10 mM EGTA, 50 mM MES, 0.5 mM DTT, 6.56 mM MgCL2, 5.77 mM ATP, 20 mM Imidazole and 15 mM phosphocreatine, pH 7.1). Tissue was trimmed of surrounding fat tissue and fibers mechanically separated on ice. Separated fibers were permeabilized with BIOPS solution containing 50 µg/mL saponin for 20 minutes at 4 °C. Following permeabilization, fibers were washed for 15 minutes in ice cold mitochondrial respiration solution (MIR05, 0.5 mM EGTA, 3 mM MgCl2, 60 mM K-lactobionate, 20 mM taurin, 10 mM KH2PO4, 20 mM HEPES, 110 mM sucrose and 1 g/L BSA, pH 7.1). Fibers were then blotted dry, weighed (3-5 mg total tissue weight) and placed in a Oxygraph-2K (OROBOROS Instruments) chamber containing 2 mL of 37 °C MirO5 (supplemented with 10 µM blebbistatin and 20 mM creatine). Routine oxygen consumption was measured by the sequential addition of the following substrates: malate (0.5 mM), glutamate (10 mM) and pyruvate (5 mM) to assess complex I mediated LEAK respiration. Adenosine diphosphate (ADP, 5 mM) to assess maximal complex I maximal respiration followed by succinate (10 mM) to measure OXPHOS (complex I and II mediated respiration). The uncoupling agent FCCP (carbonyl cyanide p-trifluoro-methoxyphenyl hydrazone, 0.5 µM, titrated 3X) was then added to determine maximal electron transport system (ETS) capacity. A period of stabilization followed the addition of each substrate and the oxygen flux per mass was recorded using the DatLab Software (OROBOROS Instruments).

### Mitochondrial content

DNA was isolated from 25 mg of either whole or white gastrocnemius of S1P^smKO^ and WT mice using the DNeasy Blood & Tissue Kit (Qiagen) following manufacturer’s instructions. DNA concentrations were measured via NanoDrop (Thermo Scientific) and 10ng of DNA was used for qPCR analysis using primers specific to a mitochondrial encoded gene DNA (Cox2) and nuclear encoded gene (36B4) and Power SYBR Green (Applied Biosystems). Transcripts were quantified on an ABI QuantiStudio 3 sequence detection system (Applied Biosystems) and Cox2 expression was normalized to 36B4 expression, and results analyzed using the 2^-ΔΔCt^ method and reported as relative units to controls. Primer sequences are listed in Supplemental Table 1.

### Western blotting

Skeletal muscle whole protein lysates were generated by homogenizing tissues in lysis buffer (20 mM Tris, 15 mM NaCl, 1 mM EDTA, 0.2 % NP-40, and 10 % glycerol) supplemented with 2x Protease Complete cocktail tablet (Roche) and 1x Phosphatase Inhibitors (Roche, Mannheim, Germany) with stainless steel beads in a TissueLyzer II (Qiagen). Protein lysates were rotated for 45 min at 4 °C, followed by centrifugation at 15,000 x g for 15 min at 4 ° C. Protein was quantified by bicinchoninic acid assay (BCA, Pierce Biotechnology), equal amounts of protein were resolved on a 4-15 % SDS-PAGE gradient gel (Bio-Rad), and transferred to PVDF-FL membrane (MilliporeSigma). Blots were probed with appropriate primary and secondary antibodies and proteins visualized by LI-COR Odyssey imaging system. To visualize phosphorylated Smad 2, blots were incubated with SignalFire ECL Reagent (Cell Signaling) and protein visualized with a BioRad ChemiDoc XRS+. Primary antibodies used: OXPHOS (MS604-300, Abcam); Phospho-Smad2 (Ser465/467) (3108, Cell Signaling); Smad 2 (5339, Cell Signaling); Phospho-AKT (4060, Cell Signaling), and Total-AKT (2920, Cell Signaling). Densitometry analysis was performed using LI-COR Image Studio Lite.

### Statistical Analysis

Data were analyzed using either Excel or GraphPad Prism version 9, unless indicated otherwise in Methods. A p-value <0.05 was considered statistically significant. Data are reported as ± SEM. Unpaired two-tailed Student’s t-tests were used.

## REFERENCES

1. Moore, T. M., Lin, A. J., Strumwasser, A. R., Cory, K., Whitney, K., Ho, T., Ho, T., Lee, J. L., Rucker, D. H., Nguyen, C. Q., Yackly, A., Mahata, S. K., Wanagat, J., Stiles, L., Turcotte, L. P., Crosbie, R. H., and Zhou, Z. (2020) Mitochondrial Dysfunction Is an Early Consequence of Partial or Complete Dystrophin Loss in mdx Mice. Front. Physiol. 10.3389/FPHYS.2020.00690

2. Hughes, M. C., Ramos, S. V., Turnbull, P. C., Rebalka, I. A., Cao, A., Monaco, C. M. F., Varah, N. E., Edgett, B. A., Huber, J. S., Tadi, P., Delfinis, L. J., Schlattner, U., Simpson, J. A., Hawke, T. J., and Perry, C. G. R. (2019) Early myopathy in Duchenne muscular dystrophy is associated with elevated mitochondrial H2O2 emission during impaired oxidative phosphorylation. J. Cachexia. Sarcopenia Muscle. 10, 643

3. Tezze, C., Romanello, V., Desbats, M. A., Fadini, G. P., Albiero, M., Favaro, G., Ciciliot, S., Soriano, M. E., Morbidoni, V., Cerqua, C., Loefler, S., Kern, H., Franceschi, C., Salvioli, S., Conte, M., Blaauw, B., Zampieri, S., Salviati, L., Scorrano, L., and Sandri, M. (2017) Age-Associated Loss of OPA1 in Muscle Impacts Muscle Mass, Metabolic Homeostasis, Systemic Inflammation, and Epithelial Senescence. Cell Metab. 25, 1374–1389.e6

4. Ibebunjo, C., Chick, J. M., Kendall, T., Eash, J. K., Li, C., Zhang, Y., Vickers, C., Wu, Z., Clarke, B. A., Shi, J., Cruz, J., Fournier, B., Brachat, S., Gutzwiller, S., Ma, Q., Markovits, J., Broome, M., Steinkrauss, M., Skuba, E., Galarneau, J.-R., Gygi, S. P., and Glass, D. J. (2013) Genomic and Proteomic Profiling Reveals Reduced Mitochondrial Function and Disruption of the Neuromuscular Junction Driving Rat Sarcopenia. Mol. Cell. Biol. 33, 194–212

5. McPherron, A. C., Lawler, A. M., and Lee, S. J. (1997) Regulation of skeletal muscle mass in mice by a new TGF-beta superfamily member. Nature. 387, 83–90

6. Moyer, A. L., and Wagner, K. R. (2015) Mammalian Mss51 is a Skeletal Muscle-Specific Gene Modulating Cellular Metabolism. J. Neuromuscul. Dis. 2, 371–385

7. Rovira Gonzalez, Y. I., Moyer, A. L., LeTexier, N. J., Bratti, A. D., Feng, S., Sun, C., Liu, T., Mula, J., Jha, P., Iyer, S. R., Lovering, R., O’Rourke, B., Noh, H. L., Suk, S., Kim, J. K., Essien Umanah, G. K., and Wagner, K. R. (2019) Mss51 deletion enhances muscle metabolism and glucose homeostasis in mice. JCI Insight. 10.1172/JCI.INSIGHT.122247

8. Ye, J. (2020) Transcription factors activated through RIP (regulated intramembrane proteolysis) and RAT (regulated alternative translocation). J. Biol. Chem. 295, 10271–10280

9. Sakai, J., Rawson, R. B., Espenshade, P. J., Cheng, D., Seegmiller, A. C., Goldstein, J. L., and Brown, M. S. (1998) Molecular identification of the sterol-regulated luminal protease that cleaves SREBPs and controls lipid composition of animal cells. Mol. Cell. 2, 505–514

10. Ye, J., Rawson, R. B., Komuro, R., Chen, X., Davé, U. P., Prywes, R., Brown, M. S., and Goldstein, J. L. (2000) ER stress induces cleavage of membrane-bound ATF6 by the same proteases that process SREBPs. Mol. Cell. 6, 1355–1364

11. Marschner, K., Kollmann, K., Schweizer, M., Braulke, T., and Pohl, S. (2011) A key enzyme in the biogenesis of lysosomes is a protease that regulates cholesterol metabolism. Science (80-.). 333, 87–90

12. Raggo, C., Rapin, N., Stirling, J., Gobeil, P., Smith-Windsor, E., O’Hare, P., and Misra, V. (2002) Luman, the cellular counterpart of herpes simplex virus VP16, is processed by regulated intramembrane proteolysis. Mol. Cell. Biol. 22, 5639–5649

13. Kondo, Y., Fu, J., Wang, H., Hoover, C., McDaniel, J. M., Steet, R., Patra, D., Song, J., Pollard, L., Cathey, S., Yago, T., Wiley, G., Macwana, S., Guthridge, J., McGee, S., Li, S., Griffin, C., Furukawa, K., James, J. A., Ruan, C., McEver, R. P., Wierenga, K. J., Gaffney, P. M., and Xia, L. (2018) Site-1 protease deficiency causes human skeletal dysplasia due to defective inter-organelle protein trafficking. JCI Insight. 3, e121596

14. Patra, D., Xing, X., Davies, S., Bryan, J., Franz, C., Hunziker, E. B., and Sandell, L. J. (2007) Site-1 protease is essential for endochondral bone formation in mice. J. Cell Biol. 179, 687–700

15. Patra, D., DeLassus, E., Liang, G., and Sandell, L. J. (2014) Cartilage-specific ablation of site-1 protease in mice results in the endoplasmic reticulum entrapment of type IIb procollagen and down-regulation of cholesterol and lipid homeostasis. PLoS One. 9, e105674

16. Patra, D., DeLassus, E., Hayashi, S., and Sandell, L. J. (2011) Site-1 protease is essential to growth plate maintenance and is a critical regulator of chondrocyte hypertrophic differentiation in postnatal mice. J. Biol. Chem. 286, 29227–40

17. Brandl, K., Rutschmann, S., Li, X., Du, X., Xiao, N., Schnabl, B., Brenner, D. A., and Beutler, B. (2009) Enhanced sensitivity to DSS colitis caused by a hypomorphic Mbtps1 mutation disrupting the ATF6-driven unfolded protein response. Proc. Natl. Acad. Sci. 106, 3300–3305

18. Bergeron, É., Vincent, M. J., and Nichol, S. T. (2007) Crimean-Congo Hemorrhagic Fever Virus Glycoprotein Processing by the Endoprotease SKI-1/S1P Is Critical for Virus Infectivity. J. Virol. 81, 13271–13276

19. Olmstead, A. D., Knecht, W., Lazarov, I., Dixit, S. B., and Jean, F. (2012) Human Subtilase SKI-1/S1P Is a Master Regulator of the HCV Lifecycle and a Potential Host Cell Target for Developing Indirect-Acting Antiviral Agents. PLoS Pathog. 8, e1002468

20. Kim, J. Y., Garcia-Carbonell, R., Yamachika, S., Zhao, P., Dhar, D., Loomba, R., Kaufman, R. J., Saltiel, A. R., and Karin, M. (2018) ER Stress Drives Lipogenesis and Steatohepatitis via Caspase-2 Activation of S1P. Cell. 175, 133–145.e15

21. Schweitzer, G. G., Gan, C., Bucelli, R. C., Wegner, D., Schmidt, R. E., Shinawi, M., Finck, B. N., and Brookheart, R. T. (2019) A mutation in Site-1 Protease is associated with a complex phenotype that includes episodic hyperCKemia and focal myoedema. Mol. Genet. Genomic Med. 7, e00733

22. Yang, J., Goldstein, J. L., Hammer, R. E., Moon, Y. A., Brown, M. S., and Horton, J. D. (2001) Decreased lipid synthesis in livers of mice with disrupted Site-1 protease gene. Proc. Natl. Acad. Sci. U. S. A. 98, 13607–12

23. Espenshade, P. J., and Hughes, A. L. (2007) Regulation of sterol synthesis in eukaryotes. Annu. Rev. Genet. 41, 401–427

24. Gorski, J. P., Huffman, N. T., Vallejo, J., Brotto, L., Chittur, S. V., Breggia, A., Stern, A., Huang, J., Mo, C., Seidah, N. G., Bonewald, L., and Brotto, M. (2016) Deletion of Mbtps1 (Pcsk8, S1p, Ski-1) Gene in Osteocytes Stimulates Soleus Muscle Regeneration and Increased Size and Contractile Force with Age. J. Biol. Chem. 291, 4308–4322

25. Picard, M., Hepple, R. T., and Burelle, Y. (2012) Mitochondrial functional specialization in glycolytic and oxidative muscle fibers: Tailoring the organelle for optimal function. Am. J. Physiol. - Cell Physiol. 302, 629–641

26. Jornayvaz, F. R., and Shulman, G. I. (2010) Regulation of mitochondrial biogenesis. Essays Biochem. 47, 69–84

27. Zhang, Y. E. (2017) Non-Smad Signaling Pathways of the TGF-β Family. Cold Spring Harb. Perspect. Biol. 9, a022129

28. Derynck, R., and Budi, E. H. (2019) Specificity, versatility and control of TGF-β family signaling. Sci. Signal. 10.1126/SCISIGNAL.AAV5183

29. Guillet-Deniau, I., Mieulet, V., Lay, S. Le, Achouri, Y., Carré, D., Girard, J., Foufelle, F., and Ferré, P. (2002) Sterol regulatory element binding protein-1c expression and action in rat muscles: Insulin-like effects on the control of glycolytic and lipogenic enzymes and UCP3 gene expression. Diabetes. 51, 1722–1728

30. Rovira Gonzalez, Y. I., Moyer, A. L., LeTexier, N. J., Bratti, A. D., Feng, S., Peña, V., Sun, C., Pulcastro, H., Liu, T., Iyer, S. R., Lovering, R. M., O’Rourke, B., and Wagner, K. R. (2021) Mss51 deletion increases endurance and ameliorates histopathology in the mdx mouse model of Duchenne muscular dystrophy. FASEB J. 35, e21276

31. Grobet, L., Martin, L. J. R., Poncelet, D., Pirottin, D., Brouwers, B., Riquet, J., Schoeberlein, A., Dunner, S., Ḿnissier, F., Massabanda, J., Fries, R., Hanset, R., and Georges, M. (1997) A deletion in the bovine myostatin gene causes the double-muscled phenotype in cattle. Nat. Genet. 17, 71–74

32. Guardiola, O., Andolfi, G., Tirone, M., Iavarone, F., Brunelli, S., and Minchiotti, G. (2017) Induction of Acute Skeletal Muscle Regeneration by Cardiotoxin Injection. J. Vis. Exp. 10.3791/54515

33. Biltz, N. K., Collins, K. H., Shen, K. C., Schwartz, K., Harris, C. A., and Meyer, G. A. (2020) Infiltration of intramuscular adipose tissue impairs skeletal muscle contraction. J. Physiol. 598, 2669–2683

34. Dobin, A., Davis, C. A., Schlesinger, F., Drenkow, J., Zaleski, C., Jha, S., Batut, P., Chaisson, M., and Gingeras, T. R. (2013) STAR: ultrafast universal RNA-seq aligner. Bioinformatics. 29, 15–21

35. Liao, Y., Smyth, G. K., and Shi, W. (2014) featureCounts: an efficient general purpose program for assigning sequence reads to genomic features. Bioinformatics. 30, 923–930

36. Patro, R., Duggal, G., Love, M. I., Irizarry, R. A., and Kingsford, C. (2017) Salmon provides fast and bias-aware quantification of transcript expression. Nat. Methods. 14, 417–419

37. Wang, L., Wang, S., and Li, W. (2012) RSeQC: quality control of RNA-seq experiments. Bioinformatics. 28, 2184–2185

38. Robinson, M. D., McCarthy, D. J., and Smyth, G. K. (2010) edgeR: a Bioconductor package for differential expression analysis of digital gene expression data. Bioinformatics. 26, 139–140

39. Ritchie, M. E., Phipson, B., Wu, D., Hu, Y., Law, C. W., Shi, W., and Smyth, G. K. (2015) limma powers differential expression analyses for RNA-sequencing and microarray studies. Nucleic Acids Res. 43, e47

40. Liu, R., Holik, A. Z., Su, S., Jansz, N., Chen, K., Leong, H. S., Blewitt, M. E., Asselin-Labat, M. L., Smyth, G. K., and Ritchie, M. E. (2015) Why weight? Modelling sample and observational level variability improves power in RNA-seq analyses. Nucleic Acids Res. 10.1093/NAR/GKV412

