## Supplemental Figures and Methods for "Site-1 Protease inhibits mitochondrial metabolism by controlling the TGF-β target gene MSS51"

**Supplemental Materials**

### Supplemental Figure 1

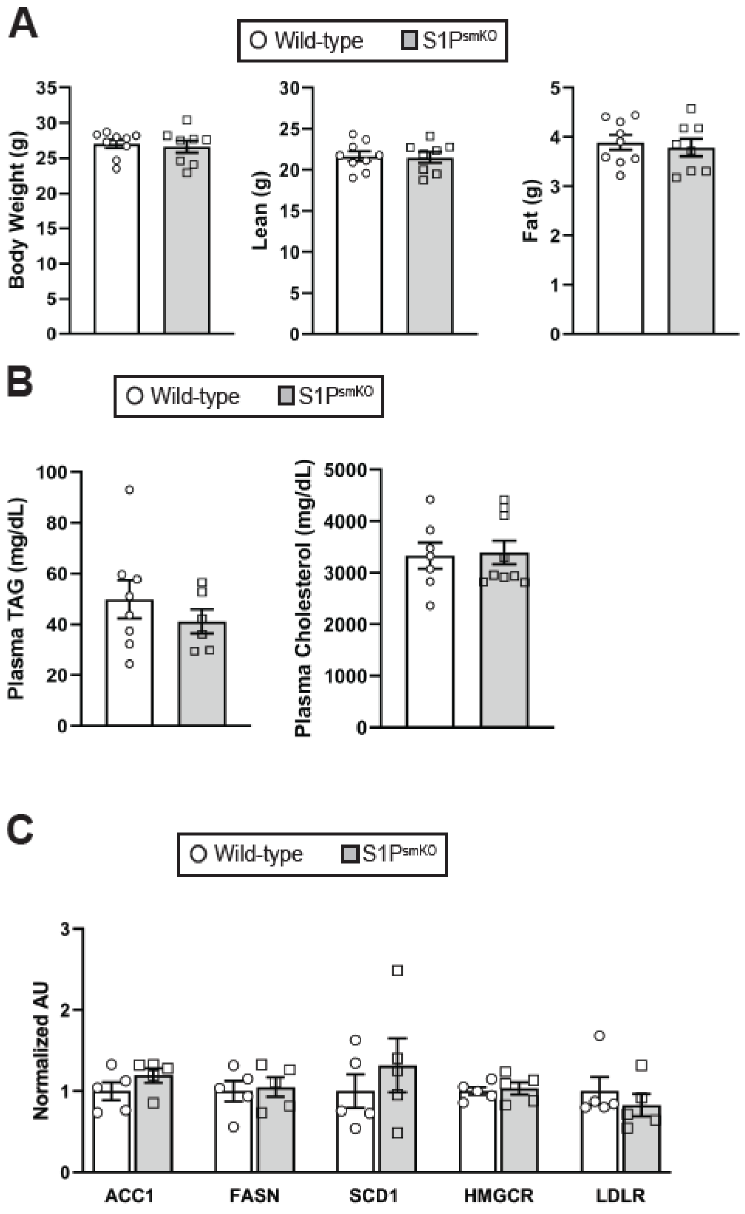


**Supplemental Figure 1. Deletion of S1P in muscle does not alter body weight or SREBP target gene expression.** (A) Body weight and body composition of S1P$^{smKO}$ and WT mice. n=7-9 per group. (B) Plasma TAG and cholesterol levels of S1P$^{smKO}$ and Wild-type mice. n=7-9 per group. (C) qPCR of SREBP target gene mRNA expression levels in gastrocnemius of S1P$^{smKO}$ and WT mice. n=5 per group. TAG, triacylglycerol.

#### Supplemental Figure 2

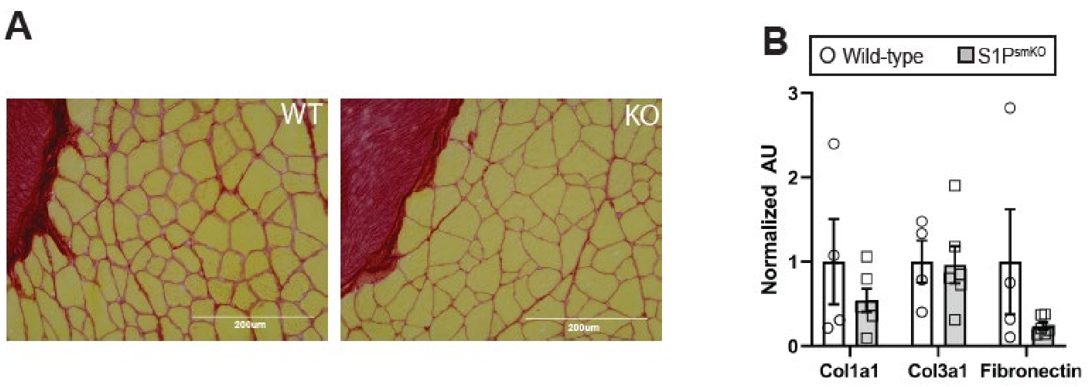


**Supplemental Figure 2. Deletion of S1P does not change age-associated muscle fibrosis.** (A) Sirius Red staining of mid-belly sections of gastrocnemius of aged S1P$^{smKO}$ and WT mice. Representative images are shown. (B) qPCR analysis of collagen and fibronectin genes in the gastrocnemius of aged S1P$^{smKO}$ and WT mice. n=4-6 per group. Gastroc, gastrocnemius; TA, tibialis anterior; WT, wild type; KO, knockout.

Primer sequences

| Gene | Forward (5'-3') | Reverse (5'-3') |
|---|---|---|
| S1P | ctg gct tct tgt gct ggt gg | ctt ttc caa agc tct cgt ccc |
| COX2 mtDNA | ctg gtg aac tac gac tgc tag a | ggc cat aga ata acc ctg gtc |
| 36B4 nDNA | acc acg aaa atc tcc aga gg | tgt cga gca ctt cag ggt ta |
| 36B4 | gca gac aac gtg ggc tcc aag cag at | ggt cct cct tgg tga aca cga agc cc |
| MSS51 | agg tct gtc cca gtt gat cct | att gga aag gcc atg agg gag |
| CHOP | cca cca cac ctg aaa gca gaa | agg tga aag gca ggg act ca |
| GRP78 | acc ccg aga aca cgg tct t | gct gca ccg aag ggt cat t |
| XBP1 | gag tcc gca gca ggt g | gtg tca gag tcc atg gga |
| ACC1 | atg ggc gga atg gtc tct ttc | tgg gga cct tgt ctt cat cat |
| FASN | gtc tgg aaa gct gaa gga tct c | tgc ctc tga acc act cac ac |
| SCD1 | ttc ttg cga tac act ctg gtg c | cgg gat tga atg ttc ttg tcg t |
| HMGCR | cca cgc agc aaa cat tgt ca | gca ggc ttg ctg agg tag aa |
| LDLR | acc tgc cga cct gat gaa ttc | gca gtc atg ttc acg gtc aca |
